# Effects of neuromuscular priming with spinal cord transcutaneous stimulation on lower limb motor performance in young active males

**DOI:** 10.1101/2025.03.09.642184

**Authors:** Simone Zaccaron, Lara Mari, Mattia D’Alleva, Jacopo Stafuzza, Maria Parpinel, Stefano Lazzer, Enrico Rejc

## Abstract

This study investigated the effects of non-invasive spinal cord transcutaneous stimulation (scTS) applied during an exercise-based priming protocol on the neuromuscular performance of lower limbs in young active males. Twelve volunteers (age: 22.7±2.1 years) participated in this randomized crossover, sham-controlled study. Maximal voluntary contraction and low-level torque steadiness of knee extensors as well as maximal explosive extension of lower limbs were assessed before and after the priming protocol with scTS or sham stimulation over a total of four experimental sessions. Further, characteristics of evoked potentials to scTS related to spinal circuitry excitability were assessed in supine position before and after the scTS priming protocol. The exercise component of the ∼25-minute priming protocol consisted of low-volume, low- and high-intensity lower limb motor tasks. scTS priming protocol tended to increase or maintain maximum isometric torque during knee extension (4.7%) as well as peak force (0.2%) and rate of force development (6.0%) during explosive lower limb extensions, whereas Sham priming protocol tended to decrease them (-4.3%, -3.3% and -15.1% respectively). This resulted in significant interactions (p=0.001 to 0.018) and medium-large differences between scTS and Sham protocols. These findings were associated with meaningful trends of some neurophysiological variables. Conversely, priming protocols did not affect low-level torque steadiness. Overall, the application of scTS during the proposed priming protocol enhanced lower limb maximal performance compared to Sham priming protocol. Future studies are warranted to assess the implementation of optimized scTS and exercise-based priming protocols during training and rehabilitation programs that include high-intensity neuromuscular efforts.

**NEW & NOTEWORTHY:** Spinal cord transcutaneous stimulation applied during an exercise-based priming protocol enhanced relevant aspects of lower limb maximal performance during isometric and explosive efforts when compared to sham stimulation; this was associated with meaningful trends of some neurophysiological variables. Conversely, neither priming protocol modulated the control of low-level, non-fatiguing torque steadiness task.

## INTRODUCTION

Muscle strength and power contribute to determine the performance of a variety of human motor tasks, ranging from activities of daily life to athletic capabilities in many sports (1–4). Further, muscle strength demonstrates high predictivity for overall morbidity and mortality (5–7), and lower limb power output is a significant predictor of functional performance in ageing (8). Strength and power are defined and limited by muscle mechanics, morphological factors and neural factors (2). In particular, neural drive to the muscles, defined as the sum of spiking activities of motor neurons, plays a crucial role in determining strength and power output (2, 9, 10). For example, force output is generally limited by the inability of the nervous system to maximally drive the muscle (i.e., activation deficit) (11–13). Neural drive and voluntary activation are also altered by central fatigue (11), which leads to a reduced recruitment and firing frequency of spinal motoneurons (14, 15).

Over the past decades, non-invasive neuromodulation techniques such as Transcranial Magnetic Stimulation (TMS) and Transcranial Direct Current Stimulation (tDCS) have been explored as potential tools for enhancing neuromuscular performance (16), with the goal of modulating the sustained neural drive to motor neurons to result in increased muscle activation and motor output (17). Only a limited number of these studies were performed in athletes and outcomes remain inconclusive, with some studies showing improvements in maximal voluntary contraction (18, 19) and power output during a vertical jump (20), while others did not find significant changes (21) or even noted a deterioration in performance (22). Such discrepancy is conceivably related to differences in experimental protocol, muscle groups, participants’ characteristics and their tolerance to pain (23, 24). The positive effects of non-invasive brain stimulation on neuromuscular performance are generally linked to a depolarization of the resting membrane potential of neurons, which would mediate changes in neural excitability and lead to increased spontaneous firing rate, thereby enhancing the neural drive to the working muscle (24–27).

Electrical stimulation of the lumbosacral spinal cord is another neuromodulation approach stemming from spinal cord injury (SCI) research, and its potential to modulate neuromuscular function and performance in able-bodied individuals is still largely unknown. In the past decade, the application of invasive and non-invasive tonic spinal cord stimulation has been combined with activity-based training in individuals with SCI, promoting remarkable recovery of lower limb motor function even in those diagnosed with a chronic, clinically motor complete injury (28–32). In this population, spinal cord stimulation modulates, and often increases, the excitability of spinal circuitry so that residual supraspinal inputs crossing the lesion, which are not functional under normal circumstances (i.e., chronic paralysis), demonstrate an augmented effect to generate and modulate different motor tasks (28, 30, 31, 33).

In able-bodied healthy individuals, exploratory studies that have investigated the effects of non-invasive spinal cord stimulation on motor control and neuromuscular performance are rather scanty. For instance, the effect of transcutaneous spinal direct current stimulation or sham stimulation applied in supine position was tested during series of subsequent countermovement jumps performed up to 3 hours after the application of spinal stimulation in two separate sessions (34). Time course of the kinetic outcomes collected in this pilot study throughout the 3-hour protocol suggested opposite trends between spinal stimulation (i.e., positive trends) and sham stimulation (negative trends). Transcutaneous spinal direct current stimulation applied during a single session of backward locomotor training also appeared to improve learning and retention of such locomotor skill compared to backward locomotor training with sham stimulation (35). Spinal cord transcutaneous stimulation (scTS) with 20 Hz stimulation frequency applied at midline over the lumbosacral spinal cord also appeared to interact with postural control strategies, impairing steady standing postural control (36). Overall, the outcomes of these studies suggest that non-invasive spinal cord stimulation can interact with the spinal circuitry controlling lower limbs in able-bodied subjects; however, the related mechanisms and effects on neuromuscular performance remain uncertain.

Here, our main goal was to assess whether spinal cord transcutaneous stimulation (scTS) can be a viable option to enhance maximal neuromuscular performance of lower limbs in young, physically active individuals. In order to achieve this effect, we opted to apply scTS during an exercise-based protocol (i.e., ∼25 minutes of low-volume, low- and high-intensity lower limb motor tasks, (37)) so that spinal cord stimulation, afferent and supraspinal inputs to the spinal cord would be integrated to prime the nervous system. Our mechanistic hypothesis is that this approach could promote an increased excitability of the spinal circuitry, bringing the related neural network closer to activation threshold, amplifying the effect of descending neural drive during maximal efforts. Thus, we expected that the scTS priming protocol would promote enhanced lower limb force and power generation compared to the same exercise-based priming protocol performed with sham stimulation. Additionally, we assessed the effects of scTS and Sham priming protocols on a submaximal, non-fatiguing torque steadiness task to test the secondary hypothesis that neuromuscular control of a finer motor task involving low-level and non-fatiguing efforts would not be differentially modulated by the two priming protocols.

## MATERIALS AND METHODS

### Research participants

Twelve healthy and physically active young male individuals (mean ± standard deviation age: 22.7 ± 2.1 years; stature: 177.9 ± 7.2 m; body mass: 77.5 ± 10.1 kg; body mass index: 24.4 ± 2.1 kg/m^2^) were recruited at the School of Sport Sciences (University of Udine, Italy) to participate in this study. Research participants were non-professional athletes who practiced team sports (soccer and basketball), individual sports (mixed martial arts, fencing) and/or related conditioning training activities for 3 ± 1 days/week. Subjects had no history of neurological and orthopaedic injuries. The experimental protocol was conducted in accordance with the Declaration of Helsinki and was approved by the Institutional Review Boards of the University of Udine (IRB# 197/2023). Before the start of the study, subjects were carefully informed about its purpose and risks, and written informed consent was obtained from all of them.

### Experimental protocol

The experimental protocol of this randomized crossover, sham-controlled study was comprised of six visits to the laboratory (Fig. 1). Each experimental session lasted approximately between 1 hour (Session 6) and 1 hour 40 minutes (Sessions 2-5), and the time interval between consecutive sessions ranged between 2 to 5 days.

**Figure 1.**
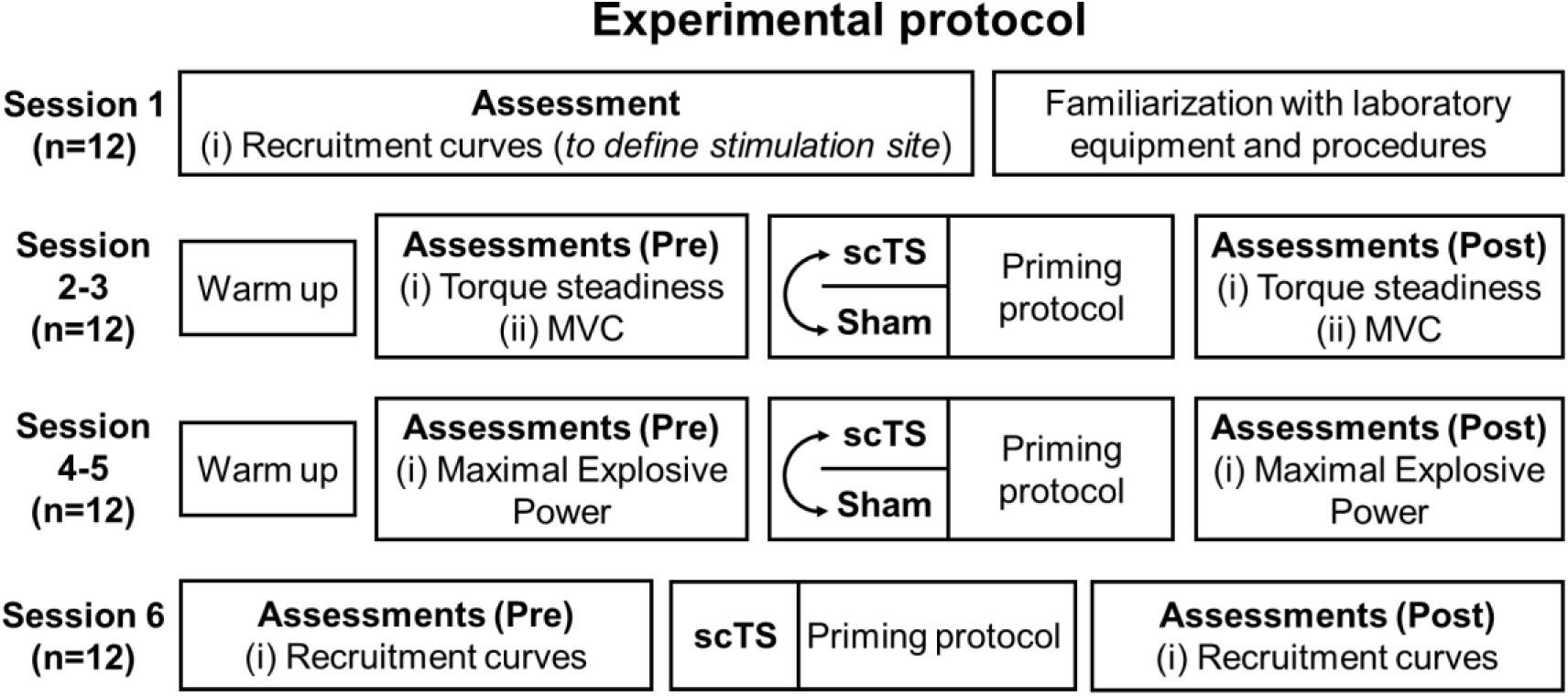
Experimental protocol. Overview of the six experimental sessions of the study. All subjects enrolled (n = 12) completed the experimental protocol. Testing of scTS or Sham priming protocol in Sessions 2-3 and 4-5 was proposed in a randomized order. MVC: maximal voluntary contraction; scTS: spinal cord transcutaneous stimulation; Sham: sham stimulation.

The first experimental session was devoted to (*i*) the assessment of anthropometric characteristics; (*ii*) determining the preferred stimulation site by assessing the relationship between scTS intensity and spinal cord-evoked potentials characteristics of lower limb muscles (i.e., recruitment curves); and (*iii*) the familiarization of participants with the laboratory equipment and procedures of the study, such as the application of scTS, isometric knee extensions on the dedicated ergometer, and explosive lower limb extensions on a sled ergometer. The second and third sessions were devoted to assessing the effects of the priming protocol with scTS or sham stimulation (in a randomized order) on maximal voluntary isometric contractions (MVC) and low-level isometric torque steadiness during knee extension. Similarly, during the fourth and fifth sessions we examined how the priming protocol with scTS or sham stimulation (in a randomized order) affected the maximal explosive power of lower limbs. Finally, the sixth session explored the effects of the scST priming protocol on the characteristics of the recruitment curves assessed with the participants relaxed in supine position.

#### Anthropometric characteristics

Body mass was measured to the nearest 0.1 kg using a manual weighing scale (Seca 709, Hamburg, Germany) with the subject wearing only light underwear and no shoes. Stature was measured to the nearest 0.5 cm on a standardized wall-mounted height board. The dominant lower limb was considered as the limb used to kick a ball (38).

#### Data acquisition system

Electromyography (EMG) and kinetic signals as well as the onset of each stimulation pulse were collected using a dedicated acquisition system (Smart DX I, BTS Bioengineering, Milan, Italy) with a sampling rate of 1000 Hz, and the related software (Smart Motion Capture System Version: 1.10.0469, BTS Bioengineering, Milan, Italy).

#### Surface EMG recordings

Surface EMG was collected using a wireless EMG system (BTS FREEEMG1000, BTS Bioengineering Milan, Italy; Input impedance: 100 MΩ; Common Mode Rejection Ratio: >110 dB @50-60 Hz; Sensitivity: 1 µV) and pre-gelled surface electrodes (BlueSensor N-00-S/25, Ambu, Penang, Malaysia) placed with inter-electrode distance equal to 20 mm. To ensure a good electrode-skin interface, prior to the application of the electrodes, the subject’s skin was shaved, rubbed with an abrasive paste, cleaned with an alcohol solution, and dry-cleaned with gauze. EMG electrodes were fixed at the beginning of the experimental session and were not removed until the end of the session. EMG was collected from the following lower limb muscles: vastus lateralis (VL), at two-third on the line from the anterior spina iliaca superior to the lateral side of the patella; rectus femoris (RF), midway between the anterior spina iliaca superior and the superior part of the patella; medial gastrocnemius (MG), on the most prominent bulge of the muscle; and tibialis anterior (TA), at one-third on the line between the tip of the fibula and the tip of the medial malleolus. (39)

#### Recruitment curves

An experienced operator identified by palpation the spaces between the spinous processes of the thoracic (T)11 and T12, and T12 and lumbar (L)1 vertebrae, which were marked with the participant standing in an upright position. The stimulation was administered with the cathode electrode placed on the skin between either the T11-T12 or T12-L1 spinous processes, in the midline over the vertebral column, using a self-adhesive, circular electrode (diameter: 25 mm) (E-CM25, TensCare, Surrey, United Kingdom). These two stimulation sites were tested in a randomized order with a 4-minute pause in between assessments. Also, two 100 x 50 mm self-adhesive electrodes (20021, Axion GmbH, Leonberg, Germany) were placed symmetrically on the skin over the iliac crests as anodes (40). After the stimulating electrodes were secured onto the skin, the participant was placed in a supine position on a standard bedded table and remained relaxed during the assessment. Stimulation was delivered as single, 1-ms monophasic square-wave pulses every 4 s. Stimulation intensity started at 5 mA and, using 5 mA increments, it was increased up to 100 mA, or the maximum intensity that did not result in discomfort for the participant. Five stimuli were delivered at each intensity. All subjects of this study achieved 100 mA as maximum stimulation intensity. A constant current stimulator (DS7A, Digitimer, Hertfordshire, UK; maximal voltage: 400V) was controlled by a trigger box (GeMS TRIGGER BOX, EMS, Bologna, Italy) and related software (Direct USB for TRIGGER BOX, Version 1.00, EMS, Bologna, Italy) to deliver the selected stimulation protocol.

During experimental session 1, recruitment curves were assessed to define the preferred stimulation site (between T11-T12 and T12-L1 spinous processes) to be used for all subsequent experimental sessions (see *Data analysis – recruitment curves* for details). In order to maximize reproducibility of cathode electrode placement across experimental sessions, the electrode location was marked on the skin with a permanent ink pen, and pictures of the electrode were also taken. During experimental session 6, the same stimulation procedure (using only the selected stimulation site) was performed at baseline and four minutes after the completion of the priming protocol with scTS to explore its effects on neurophysiological variables related to the excitability of the spinal circuitry (Fig. 1).

#### Isometric knee extension: torque steadiness and MVC

A custom-built chair ergometer instrumented with a torque sensor for each of the two attachments was connected to the acquisition system to assess torque steadiness and MVC during knee extension. The subject was seated on the chair with his trunk secured in an upright position with a dedicated belt in order to prevent any movement. The knee angle of the left and right lower limb was set at 100 degrees by properly positioning the chair attachments, and a strap was tightened around each ankle to prevent movement. During the torque steadiness trials, real-time visual feedback depicting the real-time torque exertion and the torque target (20%MVC, based on data collected during the first experimental session) was provided on a monitor positioned in front of the subject. Participants were instructed to gradually achieve the torque target and maintain the torque signal on the torque target line with the least possible error for the subsequent 15 seconds. Two attempts for each lower limb were performed, with a 1.5-minute rest in between attempts. The subjects were subsequently asked to perform three 4-5 second MVCs for each lower limb (alternating right and left limb) as fast and forcefully as possible, with a 2-min rest between attempts.

#### Maximal explosive power

The Explosive Ergometer (EXER), described in detail by previous work from our group (41) was used to assess the maximal explosive power of the lower limbs. Briefly, the EXER consists of a metal frame supporting one rail, which was inclined by 20 degrees. A seat, fixed on a carriage was free to move on the rail, its velocity along the direction of motion being continuously recorded by a wire tachometer (LIKA SGI, Vicenza, Italy). The total moving mass of the EXER (seat and carriage together) was equal to 31.6 kg. The subject was seated on the carriage seat, secured by a safety belt tightened around the shoulders and abdomen. Two mechanical blocks were used to set the distance between the seat and the force platforms (LAUMAS PA 300, Parma, Italy), so that the knee angle at rest was 100 degrees. The blocks also prevented any countermovement during the pushing phase. Participants placed their feet sole against the force platforms in a flat standardized position, and were instructed to perform a total of four maximal bilateral lower limb extensions as fast and forcefully as possible. When the subject performed an explosive effort, he and the seat moved backwards, and the force, velocity and EMG signals were collected by the acquisition system. After each push, the subjects rested for 2 minutes with their feet placed on a dedicated support.

#### Neuromuscular priming with scTS or Sham stimulation

In the present study, an exercise-based intervention with scTS or sham stimulation, which lasted approximately 25 minutes, was implemented to prime the neuromuscular system with the goal of modulating lower limb neuromuscular performance. With the participant in standing position, stimulating electrodes were placed onto the skin over the iliac crests (anodes) and the selected intervertebral space (cathode; see *Recruitment curves* section), and were secured by elastic bandages wrapped around the participant’s trunk. A small foam piece was placed between the cathode electrode and the bandage to maintain pressure on the electrode.

Tonic, monophasic scTS was delivered at 28 Hz with 1000 µs pulse width using the constant current stimulator system described above (*Recruitment curves* section). During scST, stimulation intensity was gradually increased to achieve the highest intensity that was well tolerated by the participant in terms of comfort and ability to perform the requested motor tasks without any restriction. On average, stimulation intensity applied during the neuromuscular priming protocols with scTS was 25.4 ± 4.4 mA. scTS was well tolerated by all participants but one, who frequently reported itchy sensation in the abdomen area around the electrodes; nevertheless he completed the entire study protocol. No visible lower limb muscle contraction was elicited by scTS. On the other hand, during sham stimulation, intensity was gradually increased for only one minute, followed by a 10-second decrease and the subsequent stimulator turn off; thus, no stimulation was delivered for the remaining ∼24 minutes of the sham priming protocol (42). While we did not assess credibility and expectance of sham stimulation, three participants self-reported perception of spinal stimulation at the end of a Sham session. In particular, “Whole-body pinching sensation” and “perception of stimulation when increasing attentiveness” were reported.

The priming protocol included an initial quiet standing period with increasing scTS intensity, followed by the practice of stepping in place, unilateral balance control and unilateral quarter squats. Generally, 3 minutes of low-intensity exercise were interleaved with 30 seconds of quiet standing to complete a set; a total of three sets were performed. Finally, high-intensity attempts of the motor task tested during the session (i.e., n=3 isometric knee extensions per limb in an alternated fashion, or n=4 explosive lower limb extensions) were performed with a 2-min rest in between attempts. During experimental session 6 (i.e., recruitment curves before and after neuromuscular priming protocol), isometric knee extensions were included in the priming protocol.

### Data analysis

Data were processed using the software LabChart Reader (ADInstruments, Inc., New Zealand). EMG signals were band pass-filtered at 10-499 Hz. Torque signals collected during isometric assessments were lowpass filtered at 10 Hz. Force and velocity signals collected from the EXER during lower limb extensions were lowpass filtered at 25 Hz.

Only the best attempts (i.e., lowest coefficient of variation during torque steadiness; highest torque during MVC; highest maximal explosive power during explosive extensions) within each experimental condition (scTS or sham stimulation) and part of the session (before or after the priming protocol) were taken into consideration for further analysis.

#### Recruitment curves - Data analysis

Peak-to-peak amplitude of the evoked potentials to spinal stimulation was quantified. The average EMG peak-to-peak amplitude for each stimulation intensity (i.e., five stimuli were delivered at each stimulation intensity) was calculated. Muscle activation threshold coincided with the lowest stimulation intensity that induced reproducible evoked potentials with amplitude higher than the mean baseline EMG plus three times its standard deviation (43). The largest EMG peak-to-peak amplitude was also considered for analysis.

For each participant, the stimulation site (i.e., T11-T12 or T12-L1 spinous processes) favouring overall lower activation threshold and larger EMG peak-to-peak amplitude for the right and left VL and MG was selected and used for all subsequent sessions. For all participants, T12-L1 was selected as stimulation site. When assessing the effects of the priming protocol on the characteristics of the evoked potentials to spinal stimulation (experimental session 6), the right and left VL were considered for analysis, and EMG peak-to-peak amplitude was expressed as percentage of the largest amplitude generated before (Pre) or after (Post) the priming protocol within each muscle and research participant. Data deriving from the left and right VL were averaged and then considered for the Post *vs* Pre priming protocol analysis.

#### Torque steadiness – data analysis

The variability of torque generated during low-level (20%MVC) isometric knee extension was assessed by calculating its coefficient of variation (standard deviation/mean) over a 10-second sliding window that considered the ∼20-second attempt. The time window returning the lowest coefficient of variation was also used to assess the EMG amplitude (by root mean square) as well as median power frequency for the knee extensors considered in this study (VL and RF). EMG amplitude was expressed as percent of the MVC obtained within the same experimental session. EMG amplitude and median power frequency of VL and RF were averaged to assess the overall behaviour of knee extensors (KE) (44, 45). Data deriving from the right and left lower limb were finally averaged to assess the overall effects of the priming protocol with scTS or sham stimulation.

#### MVC – Data analysis

The maximum torque generated during isometric knee extension was defined by a 1-second sliding window. EMG amplitude and median power frequency of VL and RF were also calculated within the 1-second window corresponding to maximum torque output. Rate of torque development was characterized by its peak slope detected during torque rising. EMG amplitude was expressed as percent of the MVC obtained for each muscle and research participant within the same experimental session. EMG amplitude and median power frequency of VL and RF were averaged to assess the overall behaviour of KE. Data deriving from the right and left lower limb were finally averaged to assess the overall effects of the priming protocol with scTS or sham stimulation.

#### Maximal explosive power – Data analysis

Mechanical power developed during the maximal bilateral explosive extensions on the EXER was obtained from the instantaneous product of the bilateral forces multiplied by the backward velocity. Peak force, rate of force development, velocity and power were considered for analysis. EMG amplitude of VL, RF, MG and TA of the dominant lower limb was characterized by root mean square during the push phase (i.e. throughout the period of force development), and expressed as percent of isometric MVC performed at the beginning of the experimental session. EMG amplitude of VL and RF were averaged to assess the overall behaviour of KE.

#### Statistical analysis

Statistical analysis was performed using JASP 0.19 (Amsterdam, The Netherlands). A p value less than 0.05 was considered statistically significant. Results were expressed in figures and text as mean and standard error. The Shapiro-Wilk test was used to verify the normality of distributions. Two-way within-subjects ANOVA was implemented, with “Time” (i.e., Pre *vs* Post neuromuscular priming protocol) and “Treatment” (i.e., priming protocol with scTS or sham stimulation) as repeated measures factors. Assumption of sphericity was met as factors with only two levels of repeated measures were included in the analysis. When significant differences were found, a Bonferroni post hoc test was used to determine the exact location of the differences. Also, a paired t-test or Wilcoxon test, depending on data distributions, was used to compare the Post *vs* Pre percent difference (Post-Pre-Δ%) promoted by the scTS or Sham priming protocol. Finally, effect sizes (ES) comparing Post-Pre-Δ% values obtained with the scTS or Sham priming protocol were calculated. ES values lower than 0.20 were considered negligible, between 0.20 and 0.49 small, between 0.50 and 0.79 medium, and equal or greater than 0.80 large (46).

## RESULTS

During the two experimental sessions devoted to MVC assessment, maximum isometric torque generated by knee extensors and EMG activity of VL and RF were collected Pre and Post priming protocol with scTS or sham stimulation (Fig. 2A).

**Figure 2.**
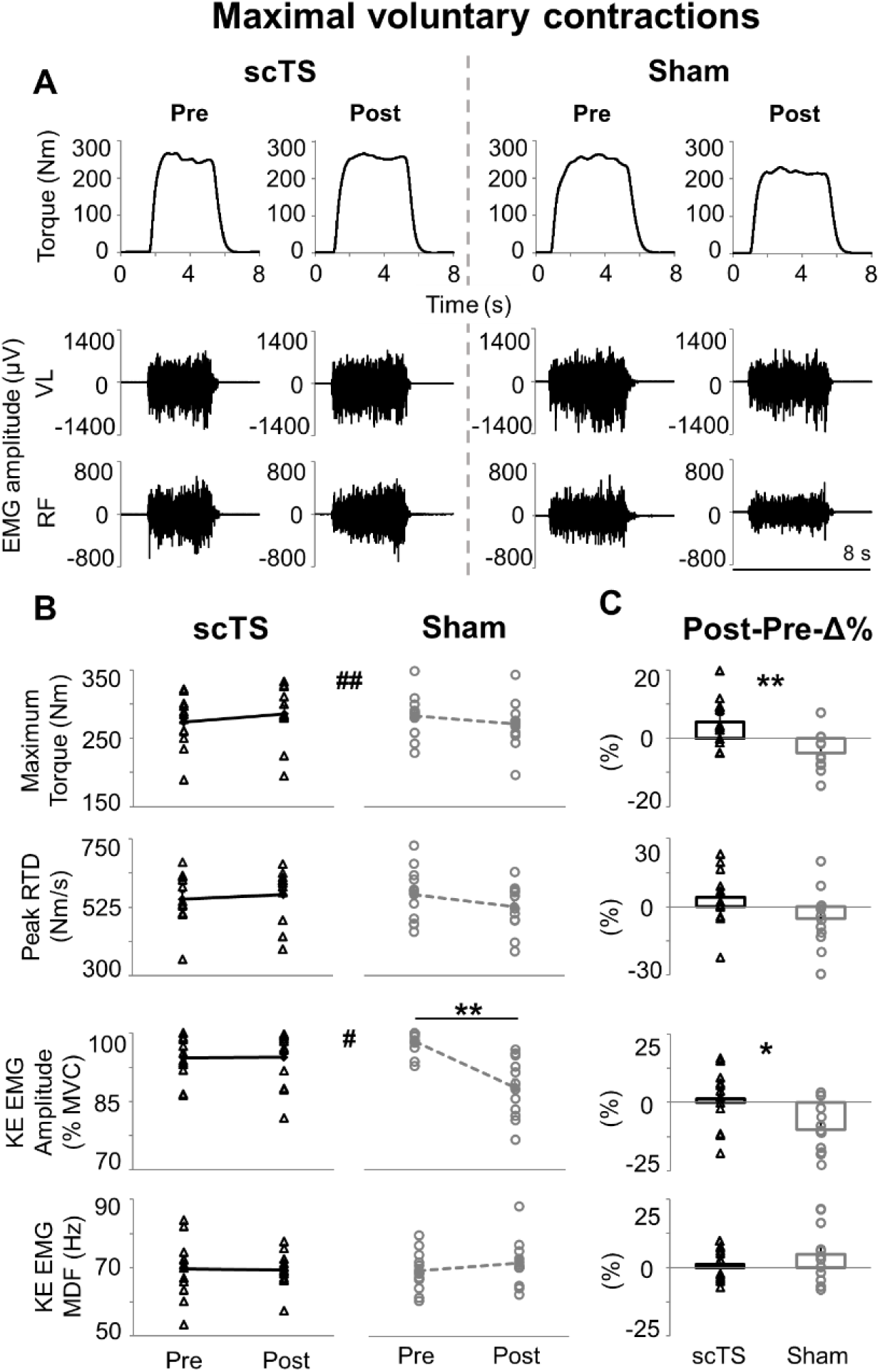
Maximal voluntary contractions. (**A**) Time course of torque output and EMG activity of vastus lateralis (VL) and rectus femoris (RF) during representative maximal voluntary isometric knee extensions generate before (Pre) and after (Post) priming protocol with spinal cord transcutaneous stimulation (scTS) or sham stimulation (Sham). (**B**) Maximum torque output, peak rate of torque development (RTD), EMG amplitude (quantified by root mean square) and median power frequency (MDF) of knee extensors (KE, average value between VL and RF). Significant Time x Treatment interaction by two-way within-subjects ANOVA: # p < 0.05; ## p < 0.01. ** significant difference by Bonferroni post hoc test, p < 0.01. (**C**) Post *vs* Pre percent difference (Post-Pre-Δ%) assessed within the scTS or Sham priming session were statistically compared by paired t-test or Wilcoxon test: * p < 0.05; ** p < 0.01. Results in (**B**) and (**C**) are reported as individual data points (empty black triangles: scTS session; empty grey circles: Sham session) as well as mean and standard error. Note that individual data points are superimposed on error bars.

No significant Treatment effect was observed for MVC-related variables, with p values ranging between 0.261 and 0.643 (KE EMG amplitude and median power frequency, respectively; Fig. 2B). Conversely, significant Time x Treatment interaction (p = 0.001) was found for the maximum torque generated, with post hoc analysis revealing a trend (p = 0.070) of lower maximum torque at Post-Sham compared to Pre-Sham. Accordingly, Post-Pre-Δ% of maximum torque was significantly different (p = 0.002, ES: 1.19; Fig. 2C) between scTS and Sham priming protocol, with a positive trend (4.67 ± 0.07%) observed after scTS priming protocol while negative values (-4.30 ± 0.06%) were found in the Sham session. A similar trend was observed for RTD (Time x Treatment interaction p = 0.062), with Post-Pre-Δ% RTD values tending to be higher (p = 0.070, ES: 0.58) in the scTS priming sessions compared to the Sham session.

These torque-related outcomes were overall supported by the EMG amplitude analysis performed on knee extensors. In particular, a significant Time x Treatment interaction (p = 0.035) was found for KE EMG amplitude, with post hoc analysis indicating lower values at Post-Sham (88.1 ± 1.9 %MVC) compared to Pre-Sham (98.1 ± 0.7 %MVC, p = 0.008; Fig. 2B) whereas similar KE EMG amplitude values (p = 1.000) were found before and after the scTS priming protocol. Along this line, Post-Pre-Δ% values of KE EMG amplitude were higher in the scTS priming session compared to the Sham session (p = 0.031, ES: 0.71; Fig. 2C), with a mean difference of 11.1% between the two sessions. Conversely, KE EMG median power frequency was not affected by the neuromuscular priming protocols herein investigated (Time x Treatment interaction p = 0.282; Post-Pre-Δ% p = 0.286, ES: 0.32; Fig. 2B and C).

A representative time course of the mechanical variables (force, velocity, power) and EMG activity of lower limb muscles during explosive efforts performed on the EXER, before and after the two neuromuscular priming protocols, is shown in Fig. 3. No significant Treatment or Time effect was observed for maximal explosive power-related variables, with p values ranging between 0.093 (KE EMG amplitude, Time effect) and 0.889 (MG EMG amplitude, Treatment effect). Conversely, a significant Time x Treatment interaction (p = 0.018) was found for the peak force, and the related Post-Pre-Δ% obtained in the scTS session was significantly higher than that assessed in the Sham session (p = 0.019, ES: 0.79) (Fig. 4). A similar finding was noted for peak RFD, which showed a significant Time x Treatment interaction (p = 0.003), and also a trend (p = 0.114) of lower values Post Sham compared to Pre Sham. Furthermore, for peak RFD, a large and significant difference between Post-Pre-Δ% obtained in the scTS and Sham session was found (p = 0.002, ES: 1.13), with higher values in the former and a mean difference of 21.1% between the two sessions (Fig. 4). No differences were observed for the other mechanical variables (peak velocity and power) considered in this assessment (Time x Treatment interaction p = 0.393 and 0.432, respectively; Fig. 4). Similarly, no significant differences, and only negligible or small effects, were noted when assessing how EMG amplitude was affected by scTS or Sham priming protocols (Fig. 4).

**Figure 3.**
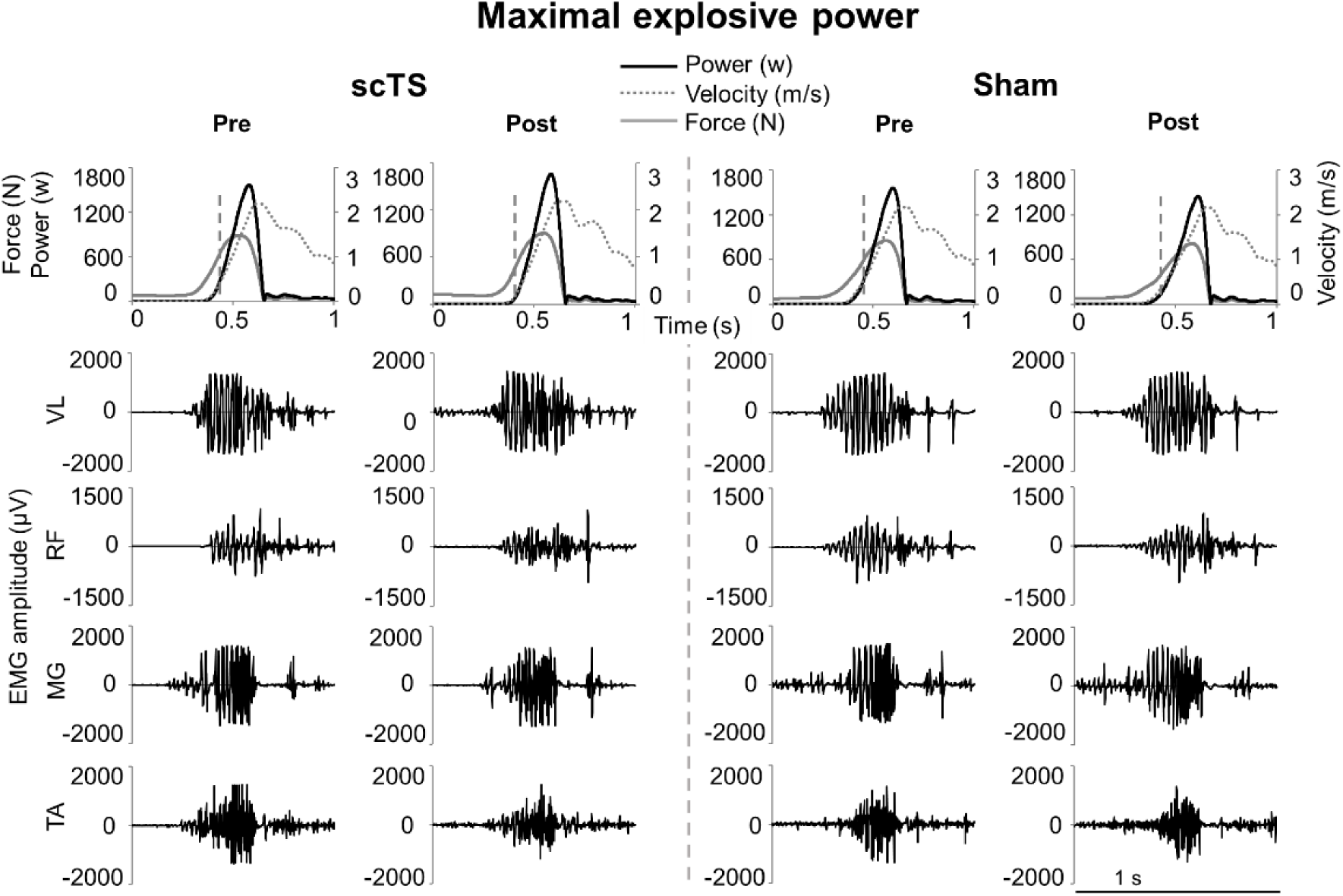
Maximal explosive power. Time course of force, velocity, power, and EMG of vastus lateralis (VL), rectus femoris (RF), medial gastrocnemius (MG) and tibialis anterior (TA) during representative bilateral explosive extensions of the lower limbs performed on the sled ergometer EXER, which was inclined by 20 degrees. Vertical, grey dashed lines in the top row indicate the time point corresponding to the peak rate of force development. These efforts were generated before (Pre) and after (Post) priming protocol with spinal cord transcutaneous stimulation (scTS) or sham stimulation (Sham).

**Figure 4.**
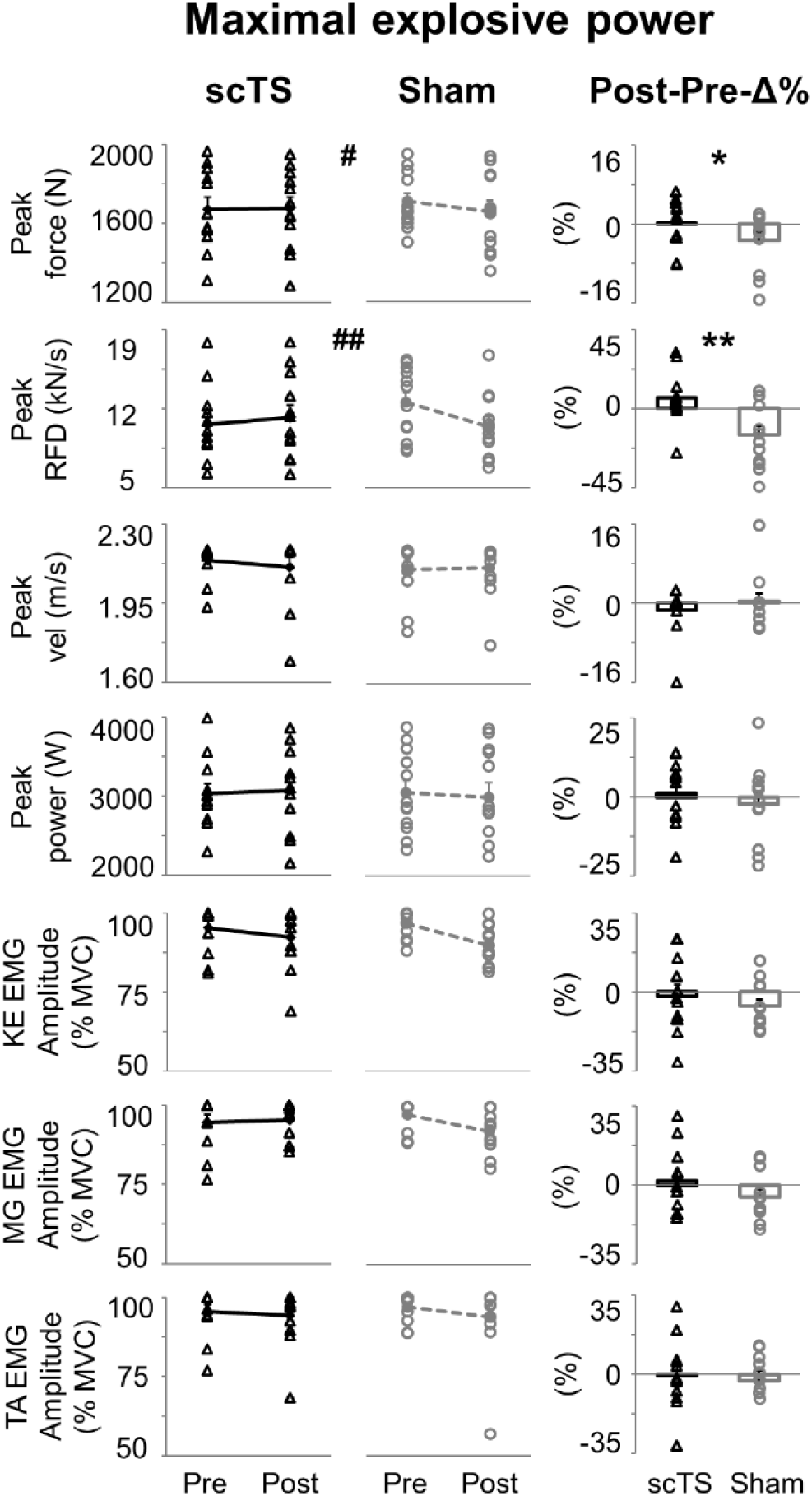
Maximal explosive power of lower limbs. Peak force, rate of force development (RFD), velocity and power as well as EMG amplitude (quantified by root mean square) of knee extensors (KE, average value between vastus lateralis and rectus femoris), medial gastrocnemius (MG) and tibialis anterior are reported as individual data points (empty black triangles: scTS session; empty grey circles: Sham session) or mean and standard error. Note that individual data points are superimposed on error bars. scTS: spinal cord transcutaneous stimulation; Sham: sham stimulation; Pre: before neuromuscular priming protocol; Post: after neuromuscular priming protocol; Post-Pre-Δ%: Post *vs* Pre percent difference within each session. Significant Time x Treatment interaction by two-way within-subjects ANOVA: # p < 0.05; ## p < 0.01. Post-Pre-Δ% calculated within the scTS or Sham priming session were statistically compared by paired t-test or Wilcoxon test: * p < 0.05; ** p < 0.01.

Effects of the neuromuscular priming protocols considered in this study on torque steadiness and EMG characteristics of knee extensors during low-level (20%MVC) efforts (Fig. 5A) were overall negligible.

**Figure 5.**
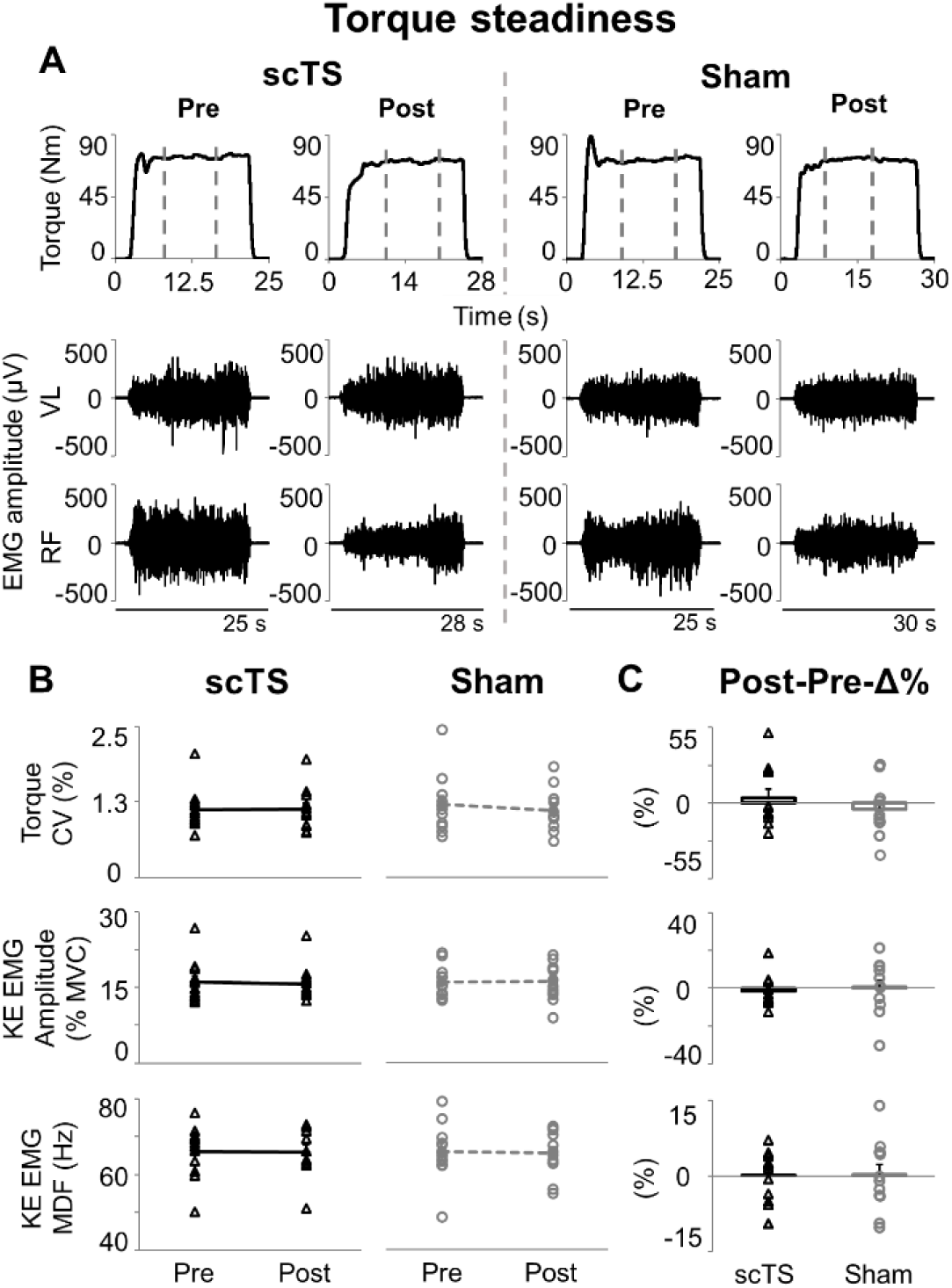
Torque steadiness. (**A**) Representative time course of torque output and EMG activity of vastus lateralis (VL) and rectus femoris (RF) during a 15-sec isometric knee extension aiming at maintaining 20%MVC torque output. Vertical grey dashed lines indicate the 10-sec time windows with the lowest torque coefficient of variation that were considered for analysis. Attempts were performed before (Pre) and after (Post) priming protocol with spinal cord transcutaneous stimulation (scTS) or sham stimulation (Sham). (**B**) Torque coefficient of variation (CV), EMG amplitude (quantified by root mean square) and median power frequency (MDF) of knee extensors (KE, average value between VL and RF). (**C**) Post *vs* Pre percent difference (Post-Pre-Δ%) assessed within the scTS or Sham priming session. Results in (**B**) and (**C**) are reported as individual data points (empty black triangles: scTS session; empty grey circles: Sham session) as well as mean and standard error. Note that individual data points are superimposed on error bars. No statistically significant difference found.

No significant Treatment or Time effect was observed for variables related to torque steadiness, with p values ranging between 0.176 (torque coefficient of variation, Time effect) and 0.824 (KE EMG amplitude, Treatment effect). Also, we found no significant Time x Treatment interactions nor meaningful trends for torque coefficient of variation (p = 0.275), EMG amplitude of KE (p = 0.393) and EMG median power frequency of KE (p = 0.857) (Fig. 5B). Additionally, no differences between Post-Pre-Δ% outcomes assessed in the scTS or Sham session were noted (torque coefficient of variation p = 0.402, ES: 0.25; KE EMG amplitude p = 0.630, ES: 0.14; KE EMG MDF p = 0.979, ES: 0.01; Fig. 5C).

Finally, characteristics of the recruitment curves explored in supine position before and after the scTS priming protocol (which included high-intensity isometric knee extensions), revealed some trends that are worth being mentioned. In particular, VL muscle activation threshold tended to be lower after scTS priming protocol (p = 0.081, ES: 0.56; Fig. 6A and B), with average stimulation intensity corresponding to activation threshold that decreased from 51.6 ± 3.2 mA to 45.5 ± 2.5 mA. Also, maximum VL EMG peak-to-peak amplitude tended to increase after scTS priming protocol (p = 0.071, ES: 0.53; Fig. 6A and C), detecting average peak-to-peak amplitude of 84.6 ± 3.9% and 95.4 ± 2.7% at Pre and Post priming protocol, respectively.

**Figure 6.**
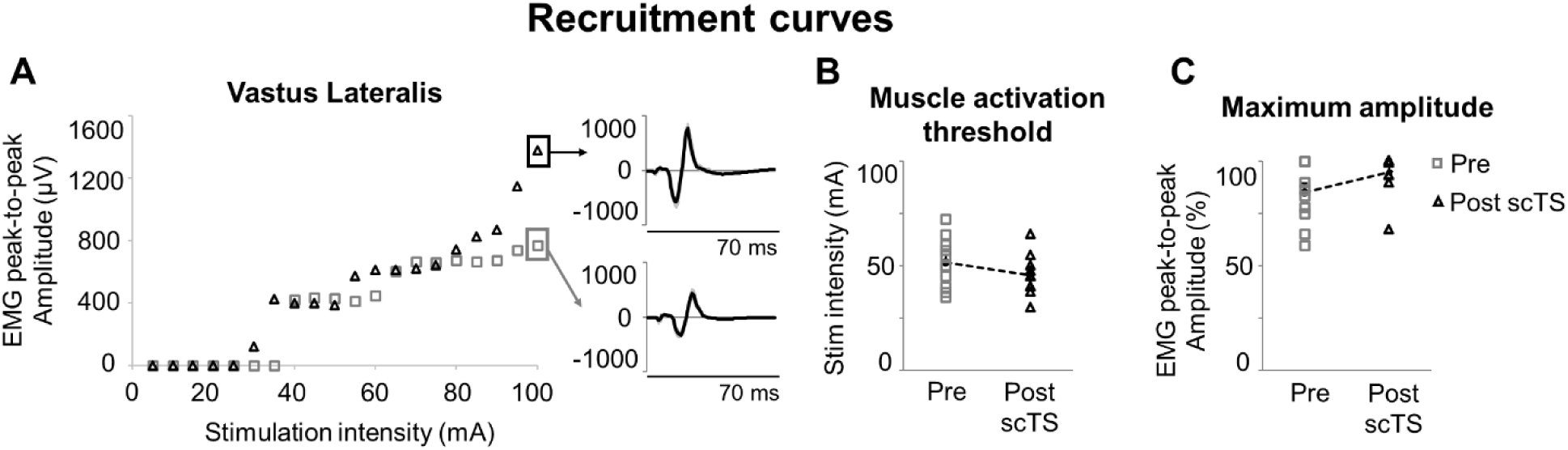
Recruitment curves. (**A**) Representative relationship between spinal cord transcutaneous stimulation (scTS) intensity and EMG peak-to-peak amplitude of vastus lateralis evoked potentials assessed before (Pre, grey empty squares) and after (Post, black empty triangles) scTS priming protocol. Each data point is the average peak-to-peak amplitude elicited by five stimuli. Exemplary evoked potentials to spinal stimulation (five individual responses are overlayed in grey; average response in black) corresponding to the maximum EMG peak-to-peak amplitude are also shown. (**B**) Vastus lateralis muscle activation threshold and (**C**) maximum EMG peak-to-peak amplitude assessed Pre and Post scTS priming protocol are reported as individual data points as well as mean and standard error. Note that individual data points are superimposed on error bars. No statistically significant difference found.

## DISCUSSION

The neuromuscular priming protocol investigated in this study differentially affected key neuromuscular variables during maximal efforts of the lower limbs depending on whether scTS or sham stimulation was applied. Generally, scTS priming protocol tended to increase mechanical outputs such as maximum isometric torque during knee extension or peak force and RFD during explosive lower limb extensions, whereas Sham priming protocol tended to decrease them. These findings, resulting in significant, medium-large differences between priming protocols, were associated with meaningful trends of some of the neurophysiological variables investigated, such as the EMG amplitude of knee extensors during maximal voluntary contractions or recruitment curve outcomes related to the excitability of the spinal circuitry. Conversely, the two priming protocols had no effect on the submaximal motor task focused on steadiness of muscle torque generation.

### Maximal neuromuscular performance of lower limbs

Descending neural drive is critical to determine maximum torque generation by recruiting the larger amount of motor units of agonist muscles at their higher firing rate and optimal synchronization (47–49). In the present study, neither neuromuscular priming protocol significantly affected maximum mechanical output within a given experimental session (i.e., Post vs Pre priming protocol, Fig. 2B). However, significant and meaningful interactions between the two priming protocols and their Post *vs* Pre effects were noted, with trends of improved performance after scTS priming, or impaired performance after Sham priming protocol (Fig. 2B and C). Negative Post *vs* Pre trends in the Sham session can be interpreted as sign of neuromuscular fatigue. In this study we did not implement neurophysiological assessments to directly examine and quantify central and peripheral components of fatigue. However, when considering the low volume of motor tasks requiring meaningful effort that were included in the priming protocol, the amount of rest between these efforts and their unilateral nature (thus doubling the rest time per leg), it is conceivable that central components primarily contributed to such negative trend. The significant decrease of KE EMG amplitude after the Sham priming protocol (Fig. 2B) further supports this perspective. Briefly, central fatigue is related to a reduced drive from the motor cortex to the muscles (50) leading to a decreased EMG amplitude and maximum force generation (47, 51). Afferent inputs deriving from muscle spindles, Golgi tendon organ as well as group III and IV fibers can also contribute to the reduction of spinal and corticospinal excitability (52).

Conversely, trends of enhanced neuromuscular performance were noted after scTS priming protocol (Fig. 2B and C). Previous research supports the view that scTS may have promoted an increased excitability of the spinal circuitry involved in control and activation of leg extensors, bringing the related neural network closer to activation threshold, thus amplifying the effect of descending neural drive during MVC (53–55). This mechanistic hypothesis originates from spinal cord injury studies showing that tonic, sub-motor threshold spinal stimulation can re-enable volitional leg movement in individuals with chronic clinically complete paralysis (29, 33, 56). In this case, the recovery of volitional movement is conceivably achieved *via* the limited spared connectivity across the injury, which carries residual neural drive that becomes sufficient to generate a muscle contraction only when the spinal circuitry distal to the injury receives appropriate electrical stimulation. Also, it is worth noting that the two priming protocols promoted differential trends in KE EMG amplitude but similar trends in KE median power frequency (Fig. 2B). This might suggest that priming protocols differentially affected the amount of motor units recruited during MVC without specificity in terms of their morpho-functional properties related to conduction velocity (57). However, future dedicated investigation involving, for example, matrix EMG and decomposition algorithms is needed to address this mechanistic hypothesis.

Some mechanical outcomes assessed during explosive lower limb extensions presented similar trends to those found during MVC. In particular, the priming protocol differentially affected peak force and RFD depending on whether scTS or sham stimulation was applied (Fig. 4), with negative Post vs Pre trends found in the Sham session and positive or no trends showed in the scTS priming session. Conversely, no significant interactions were found for peak velocity and power nor for EMG amplitude of the lower limb muscles considered for analysis. Isometric knee extension can be considered a rather simple motor task to control as it is focused on a single joint, it lacks meaningful joint movement, and its duration is relatively long (i.e., few seconds). Conversely, explosive lower limb extension performed on the EXER entails a relatively more complex motor control because it is a multi-articular, dynamic maximal effort that requires intra- and inter-muscular coordination over a very short push phase (∼300ms), which is generated against a fraction (approximately 48% in the present study) of body weight. Also, peak RFD and, to a less extent, peak force occur earlier in the push phase (i.e., corresponding to a slower instantaneous backward velocity) with respect to peak power and velocity (see Fig. 3). From these observations, we may speculate that the spinal cord excitability modulation conceivably brought about by scTS favours motor outputs related to isometric efforts and to the earlier phase of a dynamic effort. Conversely, outputs related to the later and faster phase of a dynamic effort were not supported. The investigation of RFD at different time windows in relation to force generation onset, rather than the peak RFD considered in this study, would have provided additional details on this topic. For example, RFD assessed in the early phase (0-50ms) of muscle contraction is of particular interest because it is primarily influenced by neural mechanisms (58, 59). However, a limit of the present study is that force (and torque during MVC) baseline signals were not sufficiently stable to implement a reliable detection of force generation onset to aim at such fine analysis.

### Low-level torque steadiness

Sub-maximal isometric contractions are key features of motor control, and the related steadiness of torque generation reflects the variance in descending neural drive and cumulative motor unit activity (60). In the present study, neither priming protocol affected the torque steadiness during low-level contractions (Fig. 5). EMG amplitude and median power frequency of KE were also not modulated by the priming protocols (Fig. 5). These surface EMG variables provide rather limited insights into the specific control of muscle activation. However, their lack of modulation within our study is in line with previous research detailing the relationship between force fluctuations during torque steadiness and some characteristics of the motor units recruited, such as their number, their upper limit of recruitment and their contractile properties (60). Also, it is important to consider that the motor task herein examined was non-fatiguing (i.e., 20%MVC to be maintained for approximately 15 to 20 seconds). Thus, we did not investigate aspects of motor control related to maintaining the torque target while preventing fatigue-related mechanical failure, which would intrinsically modulate the characteristics of motor units activation throughout the fatiguing contraction (61–63).

### scTS and neuromuscular priming protocol

Experimental and computational studies suggest that electrical stimulation of the spinal cord delivered with transcutaneous electrodes, or epidurally, activates common neural structures (64). In particular, spinal stimulation recruits primarily large, myelinated fibers associated with somatosensory information at their entry into the spinal cord, which in turn engages the spinal circuitry controlling muscle activation (65, 66). Parameters of spinal stimulation play a key role in determining the characteristics of activation patterns facilitation, and we leveraged previous findings deriving from spinal cord injury research to define the spinal stimulation strategy implemented in this study (30). Briefly, we initially determined for each participant the preferred cathode placement location between T11-T12 and T12-L1 spinous processes with the goal of maximizing the recruitment of both proximal and distal extensor motor pools. In fact, stimulation site has important implications for the topographical recruitment of neural structures associated with proximal and distal lower limb muscles (40). Also, we selected 28 Hz as stimulation frequency considering our goal of increasing the excitability of lower limb extensor motor pools together with a larger neural network including interneurons. Previous research suggests that stimulation frequencies between 25 and 30 Hz can engage spinal networks involved in both tonic lower limb extension and rhythmic activation patterns, and that higher frequencies progressively promote the integration of interneurons (67). Finally, stimulation intensity was always limited by participants’ comfort, and no visible lower limb muscle contraction was elicited by stimulation. This suggests that scTS neuromodulated the spinal circuitry without eliciting direct activation of motor pools.

In the present study, we explored the effects of scTS priming protocol on the characteristics of recruitment curves that can provide information on the excitability of the spinal circuitry (Fig. 6). We observed statistical trends and medium effects of scTS priming protocol on lowering muscle activation threshold and increasing maximum amplitude of evoked potentials, which are both consistent with a trend of increased excitability of the spinal circuitry. These trends, together with the main findings of the study, are also in line with previous observations of the lasting effects of spinal stimulation (i.e., after its cessation) on the spinal circuitry controlling lower limbs (34, 68, 69). It is also worth mentioning that the functional state of the spinal circuitry is dynamic and highly dependent on the afferent inputs (70). On one hand, this suggests that the interpretation of the recruitment curve findings assessed in supine position (Fig. 6) should be taken with a grain of salt when associated with the functional outcomes of the present study. On the other hand, this is the rationale that led us to apply scTS during an active neuromuscular priming protocol that provided weight bearing- and muscle contraction-related afferent inputs to the spinal cord, rather than at rest.

Briefly, the exercise-based part of the priming protocol implemented in this study was characterized by low-volume and included general low-intensity tasks (e.g., unilateral half squat and balance control) as well as task-specific, high-intensity neuromuscular components (isometric knee extension or explosive lower limbs extension), which were aimed at positively affecting neural, muscular and metabolic factors (37, 71, 72). Muscle temperature modulation and increased muscle fibre sensitivity to calcium ions are few of the potential mechanisms underlying exercise-based priming protocols. However, the priming protocol with sham stimulation tended to decrease the maximal neuromuscular performance of the lower limbs (Figs. 2 and 4), conceivably by inducing central fatigue. Previous findings indicate that performance responses to exercise-based neuromuscular priming are highly variable among individuals (73). Additionally, individuals’ performance level may influence neuromuscular potentiation and its time course, as stronger individuals may elicit greater and earlier potentiation following neuromuscular priming (74). A limit of the present study is that we did not assess the effects of Sham priming protocol on the recruitment curve characteristics. Further, it is unclear whether an optimization of the exercise-based component of the priming protocol (i.e., which would not lead to decreased performance trends with sham stimulation) would have led to larger and significant performance improvements, rather than trends, when coupled with scTS. These appear relevant topics that should be addressed in future research.

In conclusion, the application of scTS during the proposed priming protocol, which lasted approximately 25 minutes, enhanced relevant aspects of lower limb performance during maximal voluntary contraction and explosive efforts when compared to the same exercise-based priming protocol carried out with sham stimulation. Conversely, the control of a non-fatiguing submaximal motor task focused on steadiness of muscle torque generation was not affected by neither priming intervention. This suggests that scTS applied during an exercise-based priming protocol is a viable neuromodulation option to promote positive, lasting adaptations in the nervous system controlling maximal efforts of the lower limbs in healthy active males. Future studies are warranted to assess the implementation and effectiveness of optimized scTS and exercise-based priming protocols to enhance training and rehabilitation programs that include high-intensity neuromuscular efforts.

## Acknowledgments

The authors thank the study participants for their time and commitment; Teo Antognolli for contributing to data analysis. This work was supported by the Departmental Strategic Plan (PSD) of the University of Udine-Interdepartmental Project on Healthy Ageing (2020-25).

## Data availability statement

The datasets generated during and/or analyzed during the current study will be made available through material transfer agreement upon reasonable request addressed to the corresponding author.

## Author Contributions

E.R. conceived the research; E.R. and S.Z. designed the study; S.Z., L.M., M.A., J.S., M.P. and S.L. contributed to data collection; S.Z. and E.R. performed data analysis, prepared figures, interpreted the results of experiments and drafted the manuscript; all authors edited and revised the manuscript and approved its final version.

